# Olfactory and Trigeminal Routes of HSV-1 CNS Infection with Regional Microglial Heterogeneity

**DOI:** 10.1101/2024.09.22.614340

**Authors:** Christy S. Niemeyer, Laetitia Merle, Andrew N. Bubak, B. Dnate’ Baxter, Arianna Gentile Polese, Katherine Colon-Reyes, Sandy Vang, James E. Hassell, Kimberley D. Bruce, Maria A. Nagel, Diego Restrepo

## Abstract

Herpes simplex virus type 1 (HSV-1) primarily targets the oral and nasal epithelia before establishing latency in the trigeminal and other peripheral ganglia (TG). HSV-1 can also infect and go latent in the central nervous system (CNS) independent of latency in the TGs. Recent studies suggest entry to the CNS via two distinct routes: the TG-brainstem connection and olfactory nerve; however, to date, there is no characterization of brain regions targeted during HSV-1 primary infection. Furthermore, the immune response by microglia may also contribute to the heterogeneity between different brain regions. However, the response to HSV-1 by microglia has not been characterized in a region-specific manner. This study investigated the time course of HSV-1 spread within the olfactory epithelium (OE) and CNS following intranasal inoculation and the corresponding macrophage/microglial response in a C57BL/6 mouse model. We found an apical to basal spread of HSV-1 within the OE and underlying tissue accompanied by an inflammatory response of macrophages. OE Infection was followed by infection of a small subset of brain regions targeted by the TG in the brainstem, as well as other cranial nerve nuclei, including the vagus and hypoglossal nerve. Furthermore, other brain regions were positive for HSV-1 antigens, such as the locus coeruleus (LC), raphe nucleus (RaN), and hypothalamus, while sparing the hippocampus and cortex. Within each brain region, microglia activation also varied widely. These findings provide critical insights into the region-specific dissemination of HSV-1 within the CNS, elucidating potential mechanisms linking viral infection to neurological and neurodegenerative diseases.

**Importance:** This study sheds light on how herpes simplex virus type 1 (HSV-1) spreads within the brain after infecting the nasal passages. Our data reveals the distinct pattern of HSV-1 through the brain during a non-encephalitic infection. Furthermore, microglial activation was also temporally and spatially specific, with some regions of the brain having sustained microglial activation even in the absence of viral antigen. Previous reports have identified specific regions of the brain found to be positive for HSV-1 infection; however, to date, there has not been a concise investigation of the anatomical spread of HSV-1 and the regions of the brain consistently vulnerable to viral entry and spread. Understanding these region-specific differences in infection and immune response is crucial because it links HSV-1 infection to potential triggers for neurological and neurodegenerative diseases.

## Introduction

Herpes simplex virus 1 (HSV-1) is a neurotropic human double-stranded DNA virus thought to enter the central nervous system (CNS) via retrograde axon transport after infecting peripheral sensory neurons. According to the World Health Organization, herpes simplex virus 1 (HSV-1) is carried by around 66.6% of the world’s population aged from birth to 49 years old (1, 2). Notably, HSV-1 has been proposed as a trigger pathology associated with neurodegenerative diseases such as Alzheimer’s disease (AD) (3–7). Consistent with this notion, studies with human organoids also suggest that HSV-1 infection of brain tissue induces AD pathology (8). Despite the prevalence of HSV-1 and its potential impact on AD risk, little is known about which areas of the CNS are primarily infected by HSV-1 and how HSV-1 spreads throughout the CNS.

HSV-1 has the capability to infect the mucous membranes of the oral and nasal regions. It travels along the peripheral processes of trigeminal neurons, ultimately reaching the trigeminal ganglion (TG), along with other peripheral (sensory and autonomic) ganglia, where it enters a latent state (9–11). With immune challenges, such as nutritional, hormonal, psychological, or environmental stress, HSV-1 reactivates from latency, resulting in viral replication and presenting as canonical cold sores (12). This is pertinent to the risk of AD since reactivation is one of the strongest environmental drivers of AD (13) (14). In addition to latency within the TG, HSV-1 has also been shown to enter and go latent within the CNS (15, 16), and in rare cases HSV-1 can cause viral encephalitis (17).

Interestingly, HSV-1 can be found in the brain postmortem, suggesting viral entry without encephalitis (15, 18–20). The consequences of acute CNS HSV-1 infection and subsequent activation of inflammatory responses in the absence of encephalitis are unclear. While recent *in vitro* studies have shown that HSV-1 may suppress triggering receptor expressed on myeloid cells 2 (Trem2)-mediated microglial responses (21), and *in vivo* studies have noted microglial responses in thalamic regions of the brain (22), innate immune responses in multiple regions of the brain have not been described. Microglia differ in their abundance, morphology, and molecular signatures across brain regions (23, 24). Given the emerging role of microglia in neurodegenerative disease, it is pertinent to understand microglial responses in discrete sites of CNS infection by HSV-1 in the absence of encephalitis.

While HSV-1 is thought to enter the brain primarily through the monosynaptic connection between the TG and associated brainstem nuclei, HSV-1 has also been shown to enter the CNS through the olfactory route (25). The olfactory nerve is composed of axons of olfactory sensory neurons (OSNs) and provides direct access to the brain through the olfactory bulb (OB) (26). The olfactory route of infection could be relevant to cognitive problems in AD because there is a two-synapse neural connection between the OB and the hippocampus, the site of learning and memory. Herein, we aimed to characterize HSV-1 viral spread within the OE and distinct brain regions following intranasal inoculation of HSV-1. We confirmed that HSV-1 does infect distinct regions within the OE, brainstem, midbrain, and hypothalamus while sparing other regions, including the hippocampus, cortex, and OB. We also characterized the macrophage and microglial response within these distinct regions throughout the time course of acute primary infection. Together, these data provide a detailed analysis of viral entry into the brain, providing insight into HSV-1-associated CNS diseases.

## Material and Methods

### Animals

Adult C57BL/6 mice were purchased from Jackson Laboratories (Bar Harbor, ME). All animals were housed on a 14-10 light/dark cycle and fed *ad libitum* diet. All experiments outlined and animal care procedures complied with the Animal Care and Use Committee of the University of Colorado Anschutz Medical Campus.

### HSV-1 infection and tissue collection

C57BL/6 mice (male and female) are anesthetized with isoflurane followed by 20 µL of HSV-1 (10^6^ plaque-forming units/animal; McKrae strain; GenBank accession number JX142173) injected into the nares of the animal. Animals were sacrificed at 7 hours, 1-(n =3) 3-(n=2), 7-(n=3), and 10-(n=3) days post-inoculation (DPI) (**Fig. 1A-D**). In a separate cohort, male and female C57BL/6 mice were intranasally inoculated with vehicle (PBS) or HSV-1 and monitored for survival during the course of infection to 15 days post-inoculation.

**Fig. 1.**
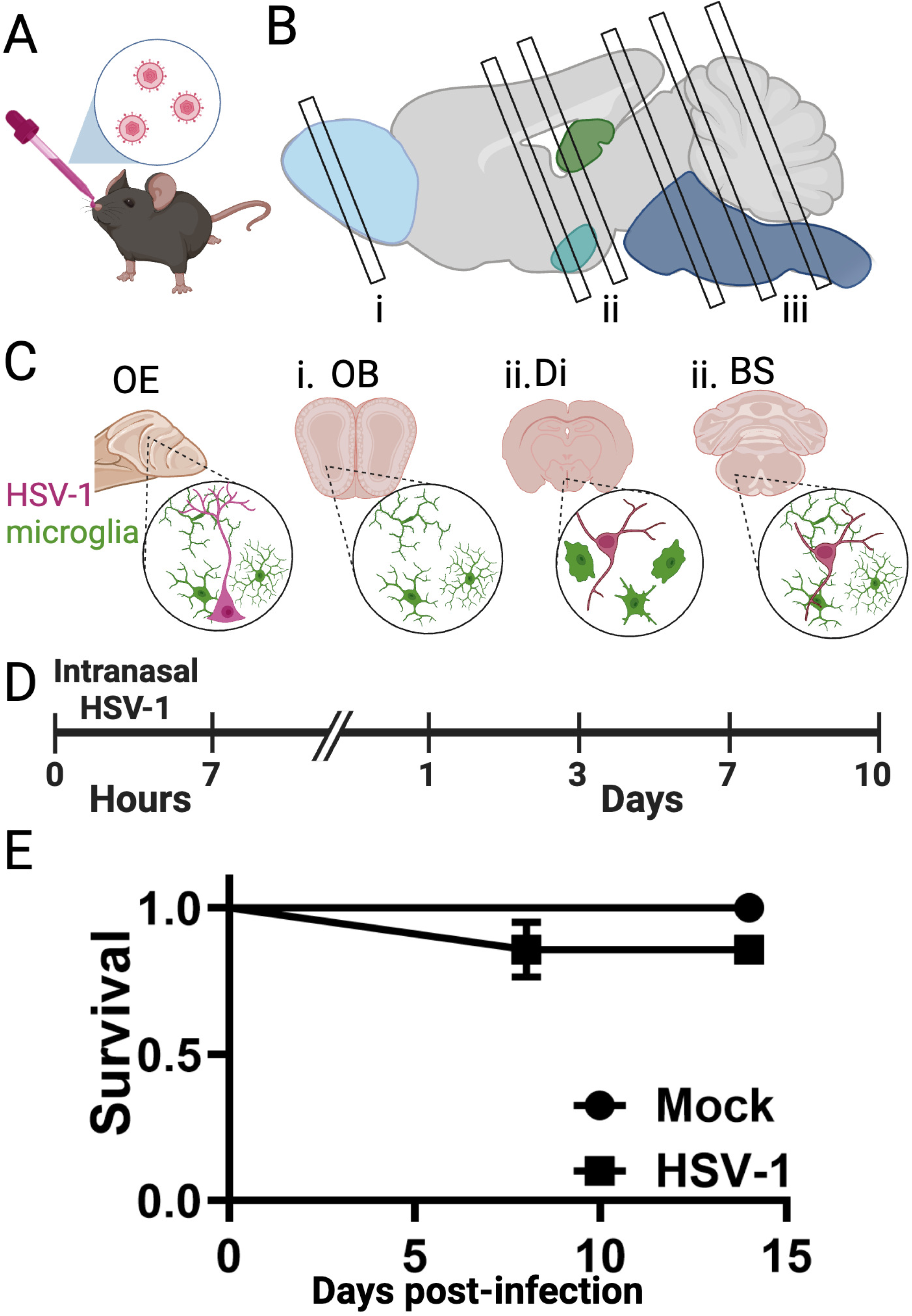
Experimental Design of intranasal inoculation of HSV-1. (**A**) C57BL/6 mice were intranasally inoculated with HSV-1 (McKrae strain, 10^6^). (**B, C**) Representation of regions of the brain is positive for HSV-1 antigen and/or microglial activation, including olfactory epithelium (OE), olfactory bulb (OB), diencephalon (Di), and brainstem (BS) subnuclei. (**D**) Timeline of infection and tissue harvest; for OE, tissue was harvested at 7 hours post-inoculation and 3 days post-inoculation with HSV-1. (**E**) For the survival curve, C57BL6 mice were intranasally inoculated with HSV-1 McKrae strain (n=14), and Mock (n=5) was followed out for 14 days post-inoculation. By 8 days post-inoculation, 2/14 HSV-1 infected animals had died but were not significantly different from controls (*p* = 0.38, by Log-rank test).

### Tissue harvest

Mice were deeply anesthetized with FatalPlus, and intracardiac perfusion was performed with ice-cold PBS 1X (∼ 15 mL) followed by ice-cold PFA 4% (∼20 mL). The nasal cavity, trigeminal ganglia, and brain were harvested for histological analysis. All samples were post-fixed in 4% PFA for 24-48h at 4°C.

Brain and TG were incubated in sucrose (30% in PBS) for 24-48h at 4°C. The nasal cavity was incubated in EDTA solution (10% in distilled water, pH 7.4) for up to one week until the bones were soft enough sectioned. The bone/OE sample was washed 3 times with PBS 1X, incubated in sucrose (20% in PBS) for 24-48h at 4°C, then embedded in OCT, frozen on dry ice, and stored at -80°C until sectioning. The nasal cavities were cryo-sectioned in the plane of the cribriform plate into 16 µm-thick sections, and sections from specific regions (**Fig. 2A**) were mounted on Superfrost Plus slides (VWR, West Chester, PA) coated with poly-D-lysine. Whole brains and TG were embedded in OCT and sliced coronally 40-45μm-thick free-floating sections mounted on Superfrost Plus slides coated with poly-D-lysine for TG. Slides were stored at -80°C, and brain-free-floating sections were stored at 4°C until use.

**Fig. 2.**
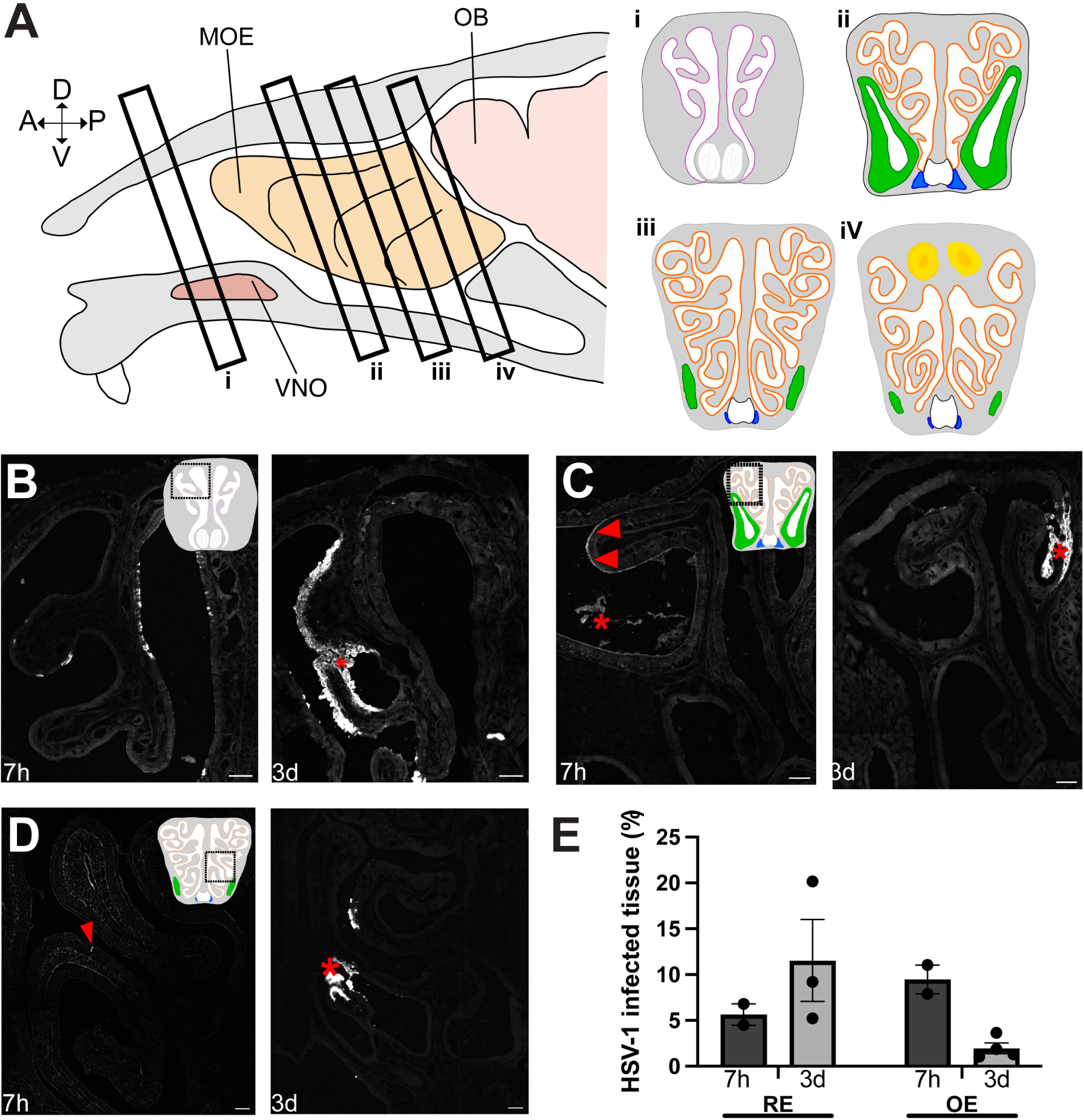
Distribution of HSV-1 antigen throughout the nasal cavity. (**A**) Diagram depicting mouse nasal structure and the specific areas where the olfactory epithelium was analyzed for immunohistochemistry. In anterior sections (**i**), the tissue lining the nasal cavity is made of respiratory epithelium (colored in magenta), whereas it is made of olfactory epithelium in posterior sections (ii, iii, and iv, colored in orange). Steno’s gland (green), nasal-associated lymphoid tissue (blue), and olfactory bulb (yellow) are also represented on the diagrams to help visualize the nasal anatomy. (**B**, **C**, **D**) Representative images of infected tissue in the nasal cavity, 7 hours, and 3 days post-inoculation with HSV-1. Debris in the lumen of the nasal cavity are indicated with an asterisk. Arrowheads point to infection of the apical layer of the olfactory epithelium. Scale bar is 100 μm. (**E**) Quantification of the surface of infected tissue across entire sections of the nasal cavity. The surface of HSV-1 positive tissue was calculated as a percentage of the total length of respiratory (RE) and olfactory epithelium (OE). 4 sections per animal were used for the quantification, one from each area described in (**A**). n=2 and n=3 animals for 7h and 3 DPI respectively. Data are represented as mean ± sem.

### Immunohistochemistry

The nasal cavity on frozen slides was dry at room temperature for 5 min, followed by incubation at 60°C for 1h. Slides were then processed as further described for brain tissue.

Free-floating brain tissue sections were incubated in heated citrate buffer (sodium citrate, pH 6.0), followed by permeabilization of the tissue with 1% Triton X-100. Brain sections were then blocked in 10% normal donkey serum. Primary antibodies were incubated overnight with 0.1% Triton X-100 and 1% normal donkey serum. Sections were incubated in secondary antibodies for two hours. Sections were then counterstained with a nuclear label (DAPI) before being mounted and were cover slipped.

**Table 1.** Antibodies used for immunohistochemistry.

| Antibody | Species | Dilution | Company (Cat#) |
| --- | --- | --- | --- |
| HSV-1 | Rabbit | 1:1000 | Dako (B011402-2) |
| IBA1 | Goat | 1:500 | Abcam (ab5076) |
| OMP | Goat | 1:500 | Wako (544-10001) |
| DAPI | N/A | 1:500 | ThermoFisher (62248) |
| Anti-rabbit 488 secondary | Donkey | 1:500 | Jackson (711-545-152) |
| Anti-goat Cy3 secondary | Donkey | 1:500 | Jackson (705-165-147) |

### Image and statistical analysis

Visualization of brain sections was performed using 3i Marianas Inverted Spinning Disk Confocal Microscope. OE’s were visualized with a Zeiss AXIO microscope, and high-magnification images of infected areas were obtained with a Leica SP8 confocal microscope. Max z-projections were generated of the region of interest within key nuclei. Images were analyzed in ImageJ.

Microglia activation was quantified branching of ionized calcium-binding adapter molecule1 (IBA1)+ cells, ranked based on activation state morphology on a scale from 1-5 (27–29): 1 for ramified microglia with long cellular processes, 2 for hyper-ramified microglia which have thicker projections, 3 for bushy microglia with a shorter ramified processes and large cell body, 4 for microglia in transitional between bushy and amoeboid states showed ‘rod like’ morphology with few short, thick processes, 5 for amoeboid microglia, with no ramified projections (**Fig. 4C**). The heatmap was generated using the pheatmap package in the statistical software program R using the mean microglia activation for each brain region at each timepoint (30).

Statistical analysis was performed using GraphPad Prism 9 (GraphPad, San Diego, CA). Statistical significance was determined using Kruskal-Wallis one-way ANOVA.

## Data Availability

All data will be made available upon request.

## Results

### HSV-1 infection with and without encephalitis in C57BL/6 mice

Survival following infection varies between HSV-1 strain, mouse genotype, and inoculation route (31, 32). Given that HSV-1 infection canonically infects humans’ oral and nasal mucosa, we examined HSV-1 spreads in the mouse following infection of the nasal mucosa by HSV-1 using a neurovirulent strain (McKrae strain). Even with a highly neurovirulent strain, the survival rate was high in C57BL/6 mice, with mortality ranging between 10-15% by 7 days post-inoculation (**Fig. 1E**). While a small number of animals had a sudden onset of sickness at days 7-8 post-inoculation, the majority of mice inoculated with HSV-1 did not present with clinical presentation of encephalitis (**Fig. S1**). We compared the spread of HSV-1 antigen at 7 DPI, and found distinct spread and accumulation of HSV-1 antigen in an animal that had succumbed to HSV-1 encephalitis compared to an HSV-1-infected animal that did not present with any clinical symptoms (**Fig. S1A, B**).

### HSV-1 infection in the nasal cavity of non-encephalitic C57BL/6

We investigated HSV-1 infection in the first potentially infected site after nasal inoculation: the nasal cavity. Immunohistochemical staining for HSV-1 revealed strong labeling in the respiratory epithelium (RE) at both 7h and 3 DPI (**Fig. 2B**). About 5% of the epithelium in the RE was already infected at 7h with HSV-1, which increased to 12% 3 DPI (**Fig 2E**). In contrast, at 7 h, HSV-1 labeling in the OE was exclusively observed in the apical edge of the OE (**Fig. 2C**, arrowhead), and HSV-1 was found sparsely within the OE layer (**Fig. 2D**, arrowhead). Cellular infected debris was also observed in the lumen of the nasal cavity near the OE-infected areas (**Fig. 2C**, asterisk). At 3 DPI, HSV-1 labeling appeared more intense in the OE, invading the epithelial layer and underlying lamina propria and was associated with cellular debris and disorganization of the laminar organization of the epithelium (**Fig. 2C, 2D**). However, in the OE these infected areas were less than 5% of the total epithelium (**Fig. 3E**). These results suggest that the viral clearance is very quick and effective in the OE, sparing most of the olfactory epithelium from infection.

**Fig. 3.**
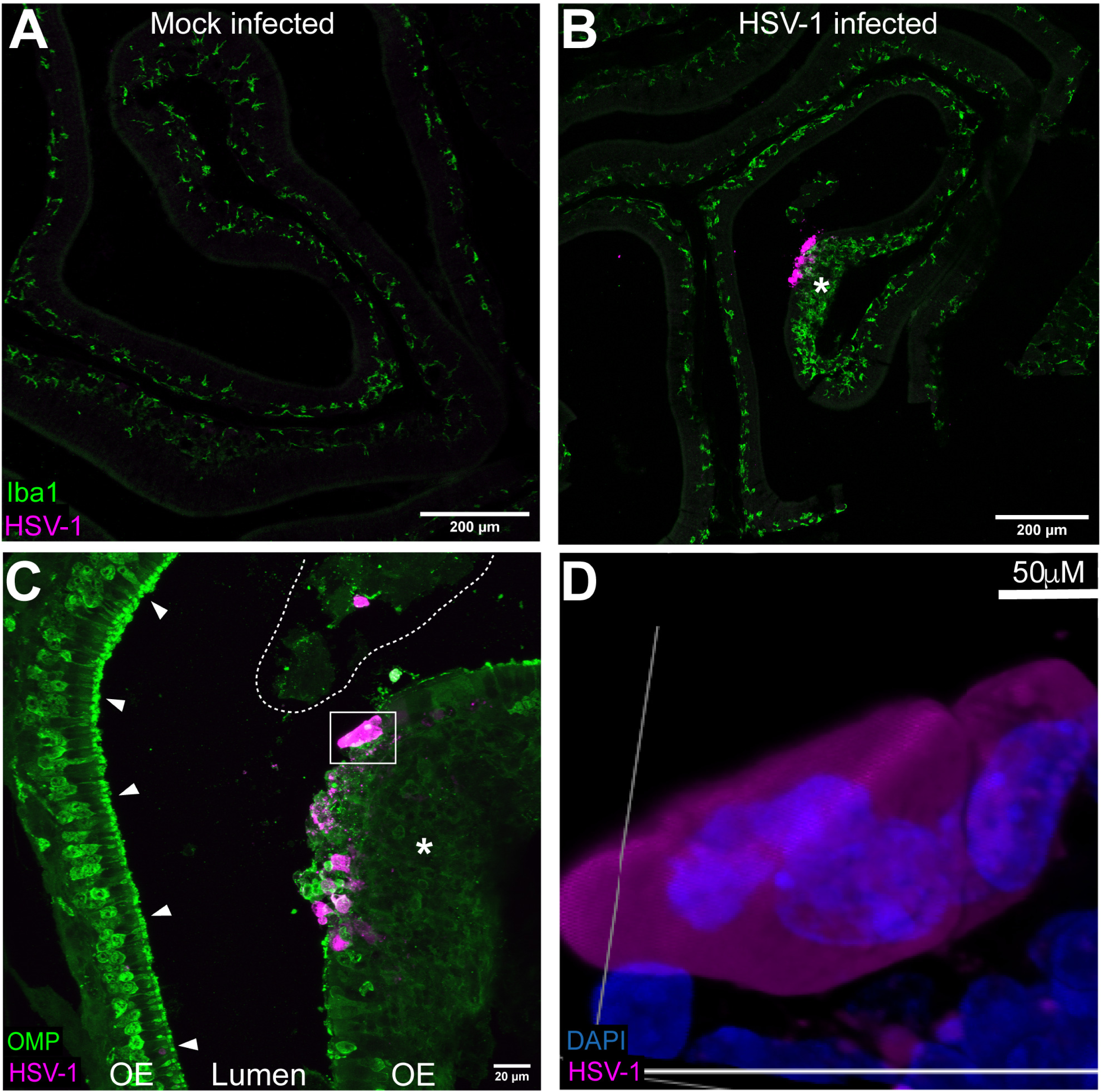
HSV-1 infection of the OE is associated with immune cell infiltration and tissue disruption. (**A**) A portion of OE from a mock-infected animal showing IBA1+ cells in the basal side of the OE. (**B**) Portion of OE from an infected animal (3 DPI) showing an infection spot in the OE (magenta) and infiltration of IBA1+ cells throughout the OE (asterisk). (**C**) Same area as in (**B**) but labeled with OMP (olfactory neurons marker). The left side (arrowheads) shows a non-infected area of OE with preserved olfactory neurons. The infection spot on the right side is associated with complete tissue disruption (asterisk). Tissue debris is also seen in the lumen of the nasal cavity (delimited with a dotted line). (**D**) High magnification image of the cell framed in (**C**) showing multiple nuclei in the same infected cell envelope.

**Fig. 4.**
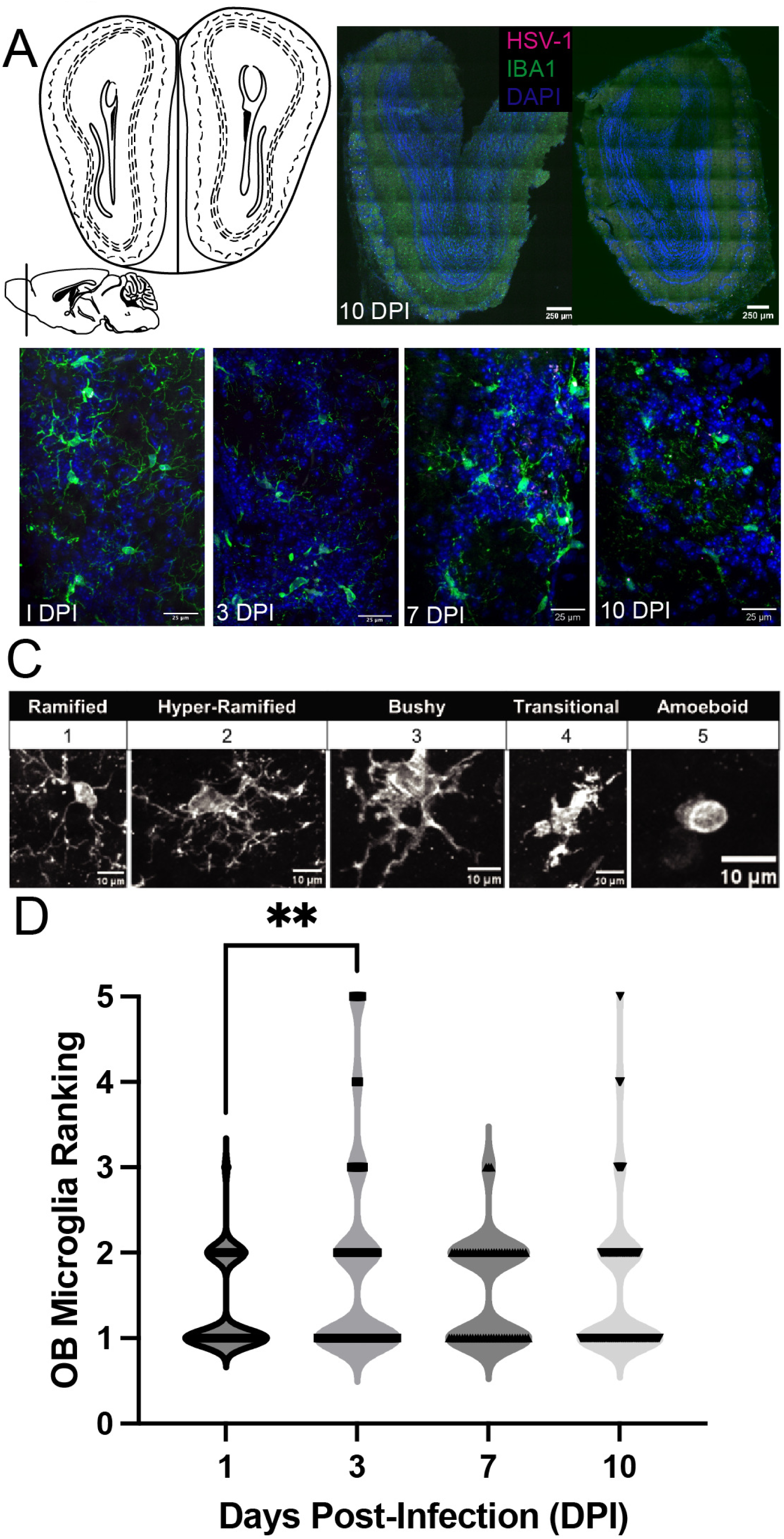
Microglia activation in the OB in the absence of HSV-1 infection. (**A**) Mouse Brain atlas image of the olfactory bulb (OB; Bregma = ∼7.32-6.56 mm) and corresponding representative histological image of a C56BL6 infected with HSV-1 at 10 days post-inoculation, stained for HSV-1 (magenta), IBA1 (green) and DAPI (blue). (**B**) Representative histological images taken from the olfactory bulb of mice inoculated with HSV-1 (Mckrae, 10^6^) at 1-,3-,7-, and 10 days post-inoculation (**C**) Schematic and representative morphology of microglial activation states based on morphology. A score of 1 = ramified microglia (small cell body and long cellular processes, 2 = hyper-ramified (thicker and more branching projections), 3 = bushy (fuzzy appearance, larger cell body, thicker projections), 4 = transitional (between busy and amoeboid, thick small projections), 5 = amoeboid= no ramified projections. **(D)** Quantification from each animal in D for microglia activation for 1 (n=188), 3 (n=144), 7(n=80), and 10(n=237) days post-inoculation. Each point is individual microglia analyzed for activation state, *p*<0.001, Kruskal-Wallis test.

We next investigated whether there was an inflammatory reaction to viral infection. We stained for IBA1, a marker for resident macrophages and dendritic cells in the OE, and OMP, a marker for mature olfactory sensory neurons (OSNs; (33, 34). As expected, in mock-infected animals, IBA1+ cells were located mostly in the lamina propria underlying the OE, with only a few cells in the basal side of the OE at 3 DPI (**Fig. 3A**). In contrast, in HSV-1-infected animals, IBA1+ cells invaded the OE, specifically near infection sites (**Fig. 3B**) that were associated with tissue disruption, as in Fig. 3C. Uninfected areas were characterized by a continuous layer of OSNs with marked OMP expression in the cell bodies and the cilia layer (**Fig. 3C**, arrowheads). Infected areas showed disrupted OMP stain with no intact OSNs (**Fig. 3C**, asterisk). The debris in the lumen of the nasal cavity was punctuated with OMP (**Fig. 3C**, delimitated), suggesting that this debris originated from the destruction of the OE. Lastly, multinucleated cells were found in the disrupted OE at the infection site, evidence of cell fusion HSV-1 induced syncytia (**Fig. 3D**).

### Time course of HSV-1 infection and microglial activation in the brain

OSN infection in the OE raises the question of whether the olfactory nerve mediates infection of the OB (35). However, we found very little HSV-1 antigen in the OB and only at 7 days post-intranasal infection compared to the other days (1-, 3-, 7, 10-DPI, **Fig. 4A**). Given there was little neuroinvasion of HSV-1 in the OB, there was still the possibility of an inflammatory response. We examined the inflammatory response by analyzing IBA1+ microglia morphology as a proxy for activation states. Morphological analysis of microglia is a common proxy for microglial activation (36, 37). We found that microglia became less ramified at 3 DPI, indicative of activation of these cells, even in the absence of HSV-1 antigen at that point (**Fig. 4C, D**).

HSV-1 is known to infect and become latent within the trigeminal ganglia, brainstem, and other cranial nerve ganglia (38). We investigated associated cranial nerve nuclei (entry points into the CNS from peripheral nerves) within the brainstem for HSV-1 antigen and found multiple sites of infection (**Fig. 5**). Brainstem nuclei associated with three cranial nerves were found infected with HSV-1: cranial nerve V (trigeminal nerve), X (vagus nerve), and XII (hypoglossal). The hypoglossal/cranial nerve XII, a purely motor nuclei involved in tongue movements, was positive for HSV-1 (XIIN, **Fig. 5A)**. Additionally, multiple vagal nerve nuclei were also positive for HSV-1, including the nucleus of the solitary tract (NoST; **Fig. 5B**), vagal nerve nuclei (DMX, **Fig. 5AC**), and area postrema (AP; **Fig. 5D).** Finally, the spinal trigeminal nucleus (SPVC) was found to be positive for HSV-1 antigen (**Fig. 5E**). The peak of infection was found throughout the brainstem at 7 DPI. However, infection appeared earlier in a few regions, including the spinal trigeminal nucleus, area postrema, and nucleus of the solitary tract at 3 DPI. No antigen was present at 10 DPI, suggesting that the virus had been cleared by that time point. We examined morphological changes of microglia at multiple time points in the brainstem (**Fig. 5**). All cranial nerve nuclei, except SPVC, had significantly activated microglia by 3 DPI. Interestingly, most cranial nerve nuclei had elevated microglial activation before the presence of HSV-1 antigen. This may suggest an inflammatory response early in infection before the presence of antigen or an early response to peripheral cranial nerve infection. We also looked at the colocalization of microglia and HSV-1 and found differences in colocalization patterns (**Fig. S2**). We found examples of microglial engulfment of HSV-1 positive cells (**Fig. S2A**). We also found examples of HSV-1 colocalizing with IBA+ (**Fig. S2B**).

**Fig. 5.**
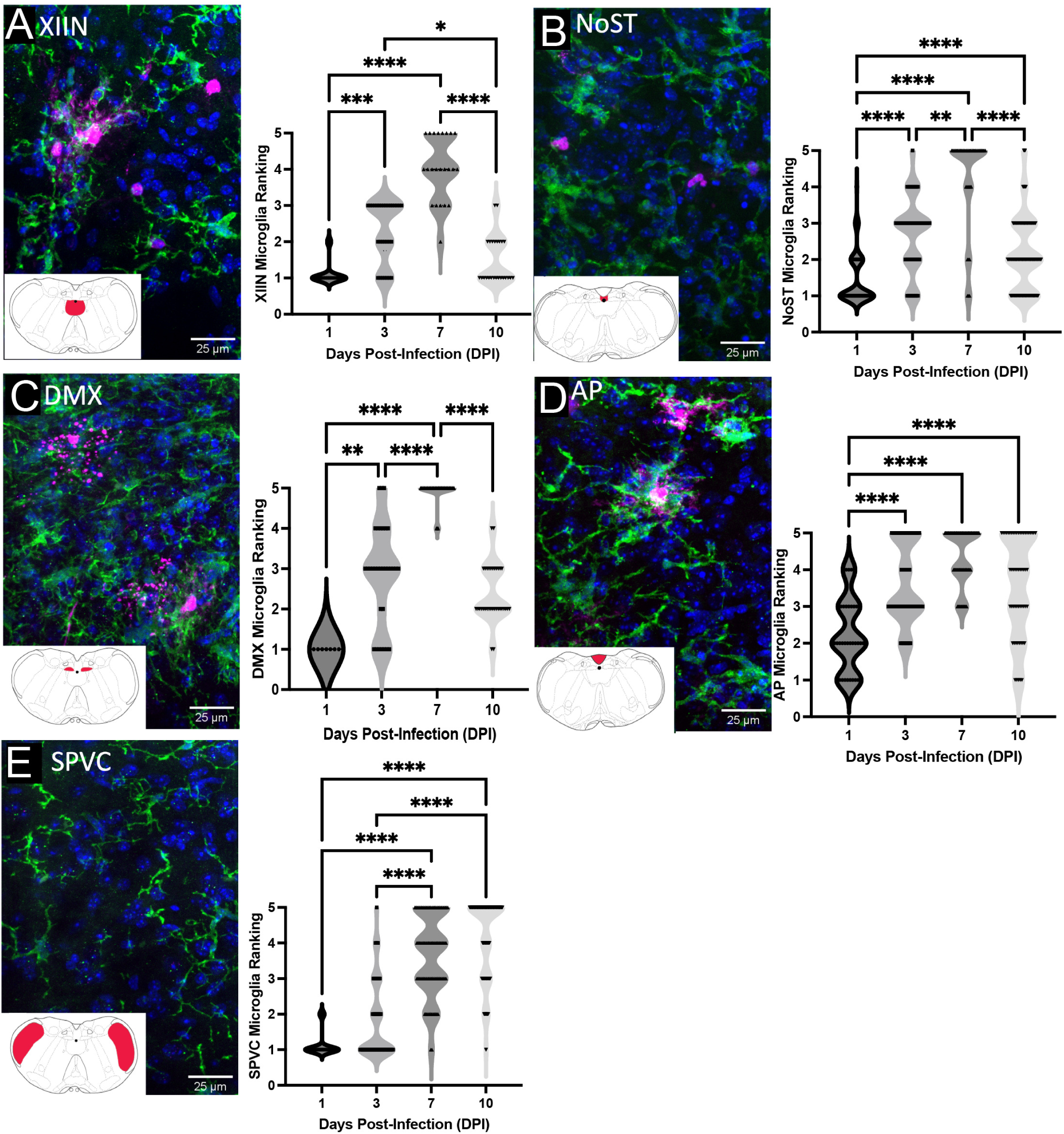
Histological assessment of microglial activation within cranial nerve brainstem nuclei. (**A**) Representative histological image of a mouse brain at 7 days post-inoculation taken from the brainstem nuclei (∼-4.04mm: -3.52mm from Bregma) cranial nerve XII nuclei (Hypoglossal nerve nuclei, XIIN). Quantification of microglia activation in XIIN at day 1 (n = 20), 3 (n = 30), day 7 (n = 23), and 10 (n = 25) post-inoculation. (**B**) the nucleus of the solitary tract (NoST). Quantification of microglia activation in NoST at day 1 (n = 90), 3 (n = 44), day 7 (n = 44), and 10 (n = 123) post-inoculation. (**C)** cranial nerve X (vagal nerve nuclei, DMX). Quantification of microglia activation in DMX at day 1 (n = 9), 3 (n = 32), day 7 (n = 39), and 10 (n = 34) post-inoculation. (**D)** area postrema (AP). Quantification of microglia activation in AP at day 1 (n = 38), 3 (n = 53), day 7 (n = 88), and 10 (n = 56) post-inoculation. (**E**) Spinal Trigeminal nerve nuclei (SPVC). Quantification of microglia activation in SPVC at day 1 (n = 54), 3 (n = 66), day 7 (n = 92), and 10 (n = 140) post-inoculation. Histological analysis of HSV-1 glycoproteins (magenta), IBA1+ microglia (green), and nuclear stain DAPI. Each point is individual microglia analyzed for activation state, *p*<0.001, Kruskal-Wallis test. * *p*<0.05, \*\**p*<0.01, *** *p*<0.001, \*\*\*\**p*<0.0001. abbreviations; XIIN, hypoglossal nucleus; DMX, vagus nerve nucleus, dorsal motor; SPVC, spinal trigeminal nucleus; NoST, nucleus of the solitary tract; AP, area postrema. Each point is individual microglia analyzed for activation state, *p*<0.001, Kruskal-Wallis test.

In addition to cranial nerve nuclei, we found HSV-1 infection in many centers of the brainstem, harboring neurons that produce neuromodulators (**Fig. 6**). Notably, we found HSV-1 labeling in primary regions of the brainstem that produce serotonin, the raphe nucleus (RaN; **Fig. 6A**). Additionally, and consistent with recent reports (16), we found HSV-1 antigen in the locus coeruleus (LC) with neurons producing norepinephrine/noradrenaline (**Fig. 6B**). We found these regions were positive for HSV-1 antigen at 7 DPI and were not labeled for HSV-1 antigen at any other time point.

**Fig. 6.**
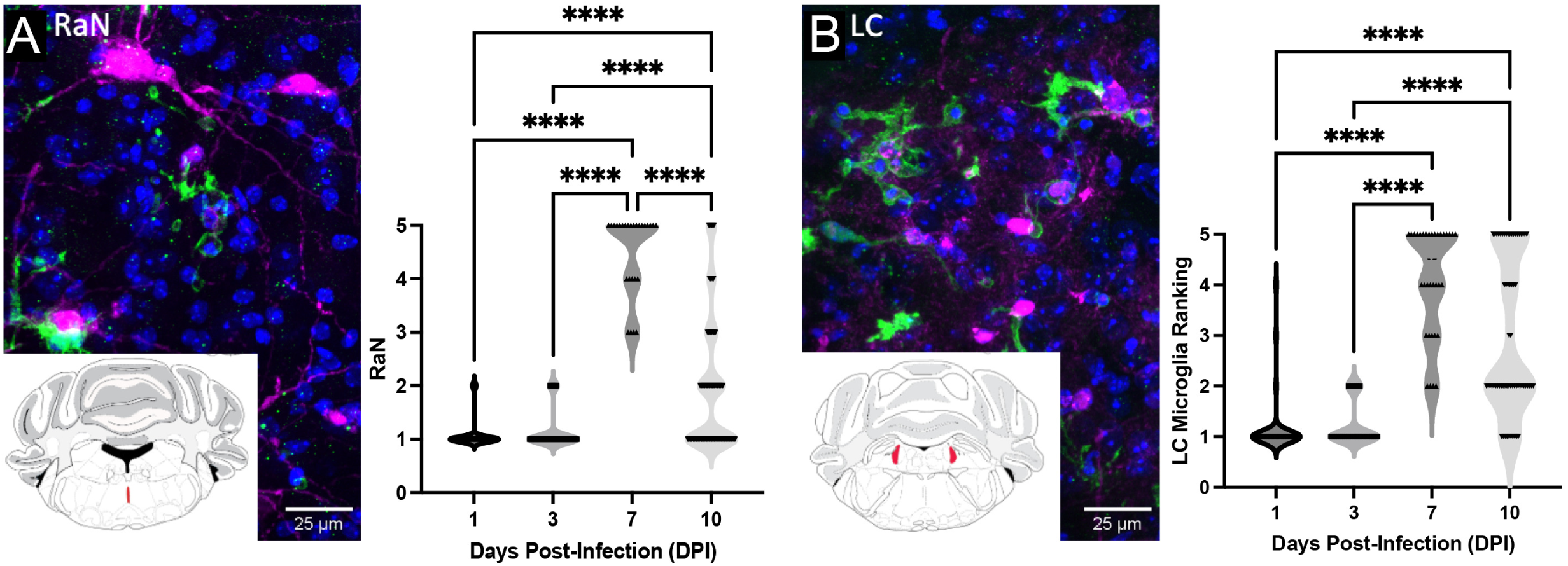
Histological assessment of microglia activation within neuromodulatory brainstem nuclei. (**A**) Representative histological image of a mouse brain at 7 days post-inoculation was taken from the brainstem nuclei (∼-3.28mm: -1.72mm from Bregma) consisting of serotonergic centers raphe nucleus (RaN). Quantification from each region of microglia activation in RaN at day 1(n = 123), 3(n = 72), day 7(n = 27), and 10(n = 132) post-inoculation. (**B**) Representative histological image of a mouse brain at 7 days post-inoculation was taken from the brainstem nuclei (∼-3.28mm: - 1.72mm from Bregma) consisting noradrenergic center the locus coeruleus (LC). Quantification from each region of microglia activation in LC at day 1 (n = 54), 3 (n = 32), day 7 (n = 34), and 10 (n = 64) post-inoculation. Each point is individual microglia analyzed for activation state, *p*<0.001, Kruskal-Wallis test. * *p*<0.05, \*\**p*<0.01, *** *p*<0.001, \*\*\*\**p*<0.0001. abbreviations; RaN raphe nucleus; LC, locus coeruleus.

Unlike the cranial nerve nuclei, where microglial activation preceded HSV-1 expression, microglial activation coincided with the presence of HSV-1 antigen at 7 DPI in neuromodulatory brainstem centers (**Fig. 6**). While microglia activation of serotonergic regions significantly decreased after viral antigen had been cleared at 10 DPI (**Fig. 6A**), noradrenergic center LC maintained an elevated microglial response at 10 DPI (**Fig. 6B**), even after HSV-1 antigen was no longer present. These results suggest a potential for elevated immune response in neuromodulatory brainstem areas, even after viral clearance.

Other nuclei were also infected within the brainstem, including several reticular formation nuclei (**Fig. S3**). Reticular nuclei have various functions, including autonomic, motor, and sensory (39). We found HSV-1 expression in the medullary reticular formation, which likely became infected because it receives trigeminal nerve branches (**Fig. S3Aii, C, E**). Medullary and intermediate reticular nuclei were also positive for HSV-1 antigen (**Fig. S3Ai, iii, B, D**). All reticular areas had a significant increase in microglia by 7 DPI, but only the medullary reticular nucleus showed sustained microglial activation at 10 DPI (**Fig. S3E-G).**

To examine the spread from the brainstem to other brain regions, we found robust infection in the subnuclei of the hypothalamus (**Fig. 7**). Specifically, we found that the lateral hypothalamus, dorsomedial nucleus, and paraventricular nucleus were positive for HSV-1 antigen at 7 DPI (**Fig. 7A-C**). Hypothalamic nuclei were positive for HSV-1 antigen by 7 DPI. Along with the brainstem HSV-1 antigen was absent by 10 DPI. Unlike brainstem regions like the LC, which had sustained microglial activation, all hypothalamic subnuclei microglial activation followed the course of infection, with significantly elevated microglia activation at 7 DPI, which went back down by 10 DPI (**Fig. 7A-C**). We did not find any HSV-1 antigen in the hippocampus, nor did we find any significantly elevated microglial activation at any time point (**Fig. 7D**). This was unsurprising, as others have found HSV-1 infection of hippocampal structures only after multiple reactivation events (6). In addition, we did not find HSV-1 antigen within the cortex (data not shown).

**Fig. 7.**
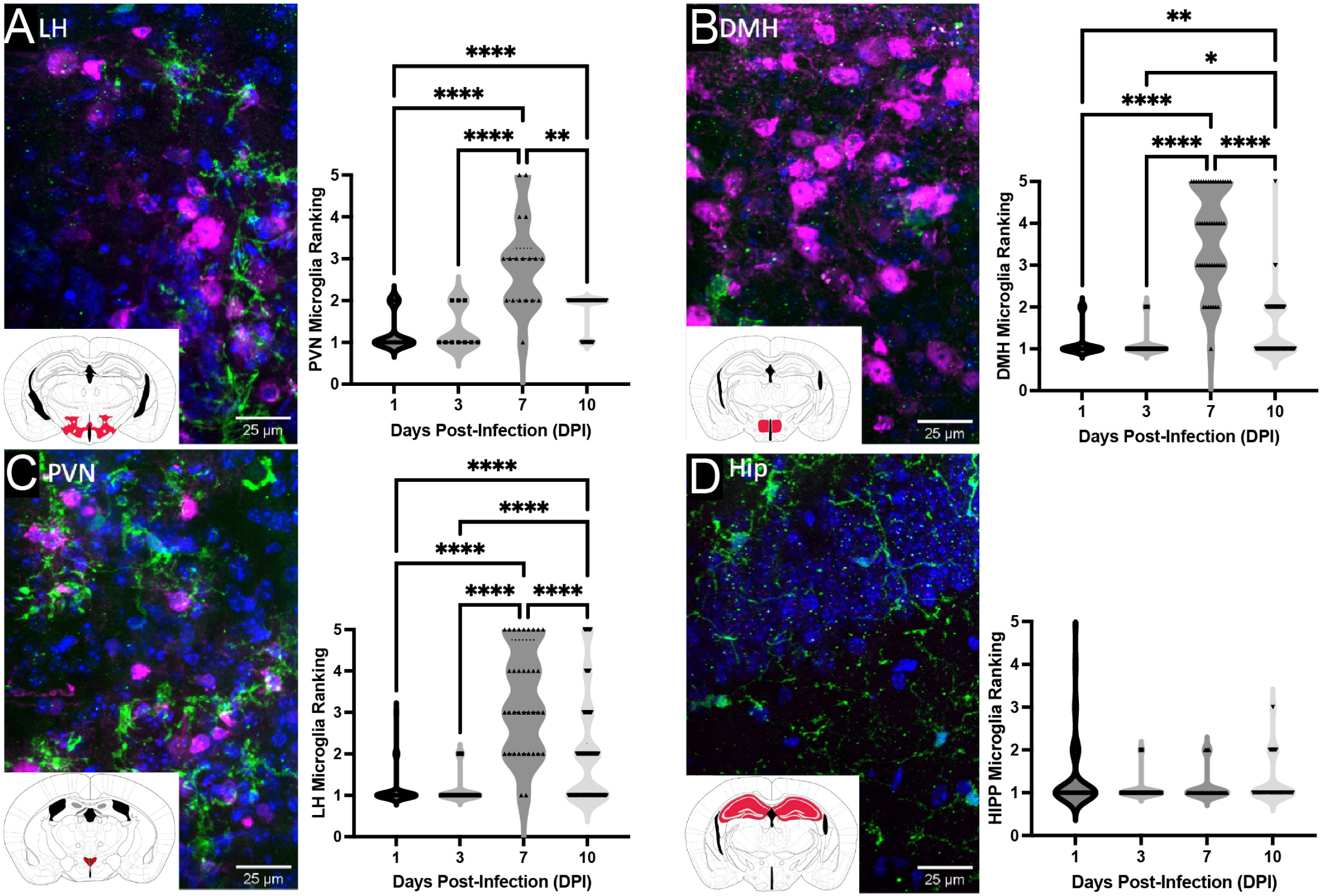
Histological assessment of microglia activation within the diencephalon sub-nuclei. (**A**) Representative histological image of a mouse brain at 7 days post-inoculation was taken through the diencephalon (∼-1.52mm: 3.08mm from Bregma) consisting of hypothalamic subnuclei, lateral hypothalamus (LH). Quantification of microglia activation in LH at day 1 (n = 136), 3 (n = 86), day 7 (n = 36), and 10 (n = 150) post-inoculation. (**B**) Representative histological image of dorsomedial hypothalamus (DMH). Quantification of microglia activation in DMH at day 1 (n = 124), 3 (n = 45), day 7 (n = 49), and 10 (n = 111) post-inoculation. (**C**) Representative histological image of paraventricular nucleus (PVN). Quantification of microglia activation in PVN at day 1 (n = 45), 3 (n = 10), day 7 (n = 18), and 10 (n = 66) post-inoculation. (**D**) Representative histological image in the hippocampus (Hip). Quantification of microglia activation in Hip at day 1 (n = 57), 3 (n = 41), day 7 (n = 5), and 10 (n = 39) post-inoculation. Each point is individual microglia analyzed for activation state, *p*<0.001, Kruskal-Wallis test. * *p*<0.05, \*\**p*<0.01, *** *p*<0.001, \*\*\*\**p*<0.0001. abbreviations; LH, lateral hypothalamus; DMH, dorsomedial hypothalamus; PVN, paraventricular nucleus; Hip, hippocampus.

### Summary of microglial activation in HSV-1 infected brain regions

We compared the activation states of microglia across all brain regions analyzed corresponding to the presence of HSV-1 antigen (**Fig. 8**). The height of activation of microglia was seen around 7 DPI, which is when we found most HSV-1 antigens (**Fig. 8A, B**). However, many regions, such as the LC, as highlighted previously, had an increase in microglia activation at 3 DPI, even though no viral antigen was found at that time. Also, it is important to highlight the OB had microglia activation at 3 DPI, which is similar to the other cranial nerve nuclei, which had early microglia activation, which may be a response to peripheral infection. We did not note microglial responses in areas such as the hippocampus. Together, these data show the distinct and heterogenic pattern of microglial response within the brain following intranasal inoculation.

**Fig. 8.**
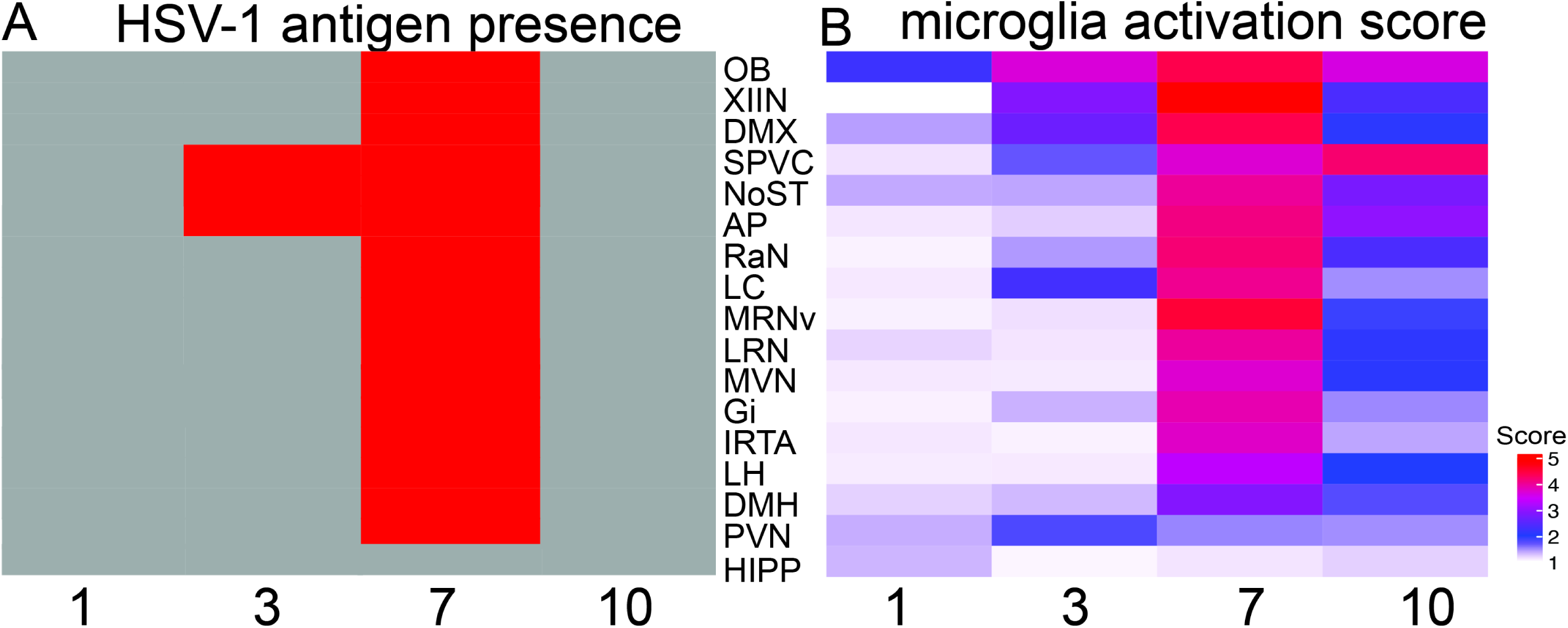
Comparison of microglial activation states within distinct brain regions over the course of HSV-1 infection. (**A**) Representation of HSV-1 antigen presence over time between each brain region. Regions that were negative (grey) or positive (red) for HSV-1 antigen are represented for each brain region. (**B**) Microglial activation states from all the above brain regions were collated and reported as an intensity measure over the course of infection (1-, 3-, 7-, and 10 days post-inoculation). Abbreviations: OB, olfactory bulb; XIIN, hypoglossal nucleus; DMX, vagus nerve nucleus, dorsal motor; SPVC, spinal trigeminal nucleus; NoST, nucleus of the solitary tract; AP, area postrema, RaN, raphe nucleus; LC, locus coeruleus, LH, lateral hypothalamus; DMH, dorsomedial hypothalamus; PVN, paraventricular nucleus; HIP, hippocampus; Gi, gigantocellular medullary reticular nucleus; MRNv, medullary nucleus-ventral; MVN, medial vestibular nucleus; LRN, lateral reticular nucleus; IRTA, intermediate reticular nucleus.

## Discussion

In the present study, we determined a heterogeneous temporal pattern of HSV-1 infection and subsequent microglia activation. Mouse brains were positive for HSV-1 antigen, and we observed microglia activation in numerous regions. Although we found a punctate infection, there was no overt sickness behavior, which was generally indicative of encephalitis, and the mice had high survival rates (**Fig. 1, Fig. S1**). We found that HSV-1 productively infects the olfactory epithelium at very early time points and only in a small fraction of the total OE (**Fig. 2**, **3**). Given the main entry into the CNS from olfactory sensory neurons in the OE, we found relatively small amounts of HSV-1 antigen in the olfactory bulb, suggesting that cranial nerve infection of the brainstem is the primary route of entry into the CNS (**Fig. 4, 5**, **Fig. S3**). We found distinct brain regions infected with HSV-1, including the brainstem’s neuromodulatory centers (serotoninergic and adrenergic) and a subset of nuclei in the hypothalamus (**Fig. 6**, **7**). Along with HSV-1 infection, we found distinct microglial activation patterns across multiple brain regions (**Fig. 8**). Temporal patterns of HSV-1 antigen expression and microglial activation were partially overlapping. The distinct regions infected with HSV-1 may underlie important physiological, pathological, and behavioral symptoms of nasal viral infection.

Our results shed light on vulnerable CNS entry sites for HSV-1 spread. Importantly, even though intranasal inoculation elicited direct infection of olfactory sensory neurons, we found very little infection in the olfactory bulb. We hypothesize that the OE mounts a rapid and robust immune response resulting in tissue disruption in order to prevent direct viral entry into the brain through OSN axons. We show HSV-1 infection was more prominent in the RE than in the OE. This observation differs from previous studies where it was suggested that HSV-1 preferentially targeted the OE as opposed to the RE (25). This discrepancy may originate from the use of different mouse and viral strains. However, our study agrees with Shivkumar *et al*. 2013 in concluding that for intranasal inoculation, HSV-1 spreads through trigeminal fibers rather than the olfactory nerve (25).

Several studies demonstrated that herpesviruses show a tropism for the OE but not the respiratory epithelium (RE) (16, 18–20). Heparan sulfate and Nectin 1 are two receptors for herpesvirus entry into cells and have been found on the apical side of the OE but are not highly expressed or are expressed on the basal side of the RE (and are thus inaccessible to the virus; (16, 18). However, despite targeting the OE, HSV-1 rarely reached the OB and did not cause significant neurological disease (16), suggesting that HSV-1 traveled along the trigeminal nerve branches instead of traveling through the olfactory nerve. The TG directly innervates the OE and OB, making it an easy access point for HSV-1 to infect the TG (40). In mouse models of HSV-1 infection, multiple sites have been shown to become productively infected and become latent (11, 21–24). Thus, our findings suggest that for intranasal infection, HSV-1 travels from the nose to the brain through the trigeminal fibers and other cranial nerves, innervating the RE and the OE (40).

It is not surprising that we found HSV-1 antigen in the spinal trigeminal nuclei of the brainstem (**Fig. 5E**). Interestingly, reticular nuclei, such as the LRN, which receive branches of the trigeminal nerve fibers, were also infected with HSV-1 (**Fig. S3Ai, B**). However, entry routes through other cranial nerves, inferred because of the expression of viral antigen in the associated brainstem nuclei, shed light on other possible entry routes that had not been identified in previous studies. The vagus nerve, in particular, was very interesting to find, as other alpha herpes viruses, such as varicella-zoster virus, have also been shown to infect the vagus nerve (41–43). Furthermore, vagal nerve disruption and associated brainstem nuclei (NoST) have long been associated with many neurological and neurodegenerative diseases, including anxiety, depression, AD, and Parkinson’s disease (44–47).

We noticed the absence of a direct correlation between HSV-1 presence and microglia activation in all regions (**Fig. 8**). In the OB, for example, microglia activation was observed at 3 DPI (**Fig. 4**), even in the absence of HSV-1 antigen expression, likely due to viral infection of the monosynapticly connected OE. Cell debris or factors released by dead cells in the OE could potentially reach the OB and act as a stress signal alerting for a potential threat (48, 49). As a result, the OB may have mounted a preventive defense mechanism through microglia activation. Microglia, immune cells, and astrocyte activation in the superficial layers of the OB have been described in numerous cases of nasal inflammation (48, 50–53). Such activation of OB immune cells may be important for removing degenerated OSN axon terminals and preventing nasal inflammation propagation to the CNS. It is also possible that HSV-1 was below the detection limit and that the microglia response was associated with early infection. Additionally, many regions infected with HSV-1 had elevated microglial response even after the virus was no longer present, including the cranial nerve nuclei SPVC, AP (**Fig. 5**), and serotonergic centers like the raphe nucleus (**Fig. 6**). Again, the asynchronous microglia activation may be due, in part, to asynchronous HSV-1 infection that was below the limit of detection for antigen but still may have been transcriptionally active. Partially overlapping temporal patterns of microglial activation and HSV-1 antigen expression document the temporal pattern of inflammatory response following HSV-1 infection.

We also examined the colocalization patterns of IBA1 and HSV-1 (**Fig. S3**). Indeed, there is debate within the literature as to whether microglia can become productively infected ( (54, 55). Taken from examples of HSV-1 antigen at the peak of infection (7DPI), we noted IBA1+ cells that were HSV-1 negative but were surrounding HSV-1 positive cells (**Fig. S2A**), indicative of microglial-engulfment of infected cells rather than productive infection. However, we also found instances of HSV-1 antigen colocalizing within IBA1+ cells (**Fig. S2B),** indicative of infection. These findings are consistent with the conflicting evidence on whether microglia themselves become infected and if that infection is productive or abortive. Further studies are needed to empirically define the dynamic microglial response to HSV-1 infection.

Microglia exhibit substantial heterogeneity across multiple domains, including distribution, phenotype, and function across the healthy brain (23, 24). The heterogeneity may arise from the developmental origins. However, interactions and influences of different microenvironments may also play a key role in shaping microglia function, such as synaptic pruning, neurogenesis, and debris clearance (56). Regional heterogeneity could underlie the selective vulnerability of brain regions to neurodegenerative diseases like AD. Microglia can play a dual role in HSV-1 infection, initially restricting viral replication via antiviral mechanisms (57, 58) but conversely also contributing to sustained neuroinflammation contributing to neurodegeneration (59). Recent reports in an HSE model found a distinct microglial response in the thalamus, contributing to an exacerbated hyperinflammatory response (22). Our results elucidate vulnerable regions of the brain that have sustained inflammatory responses after the virus has been cleared, leading to our understanding of the complex microglial response.

Many other risk factors are associated with the development of AD, including genetic variants (60–62). For example, ApoE4, an allelic variant of apolipoprotein E (ApoE), is the strongest genetic risk factor for the development of AD (63). ApoE is the major scaffold protein of brain-derived lipoproteins that transport lipids throughout the CNS (64). Although the mechanisms driving ApoE4-mediated AD pathology are still an active area of investigation, it is likely that the presence of ApoE4 results in poor lipid transport and immunometabolic disruption of microglia (65, 66). Microglial dysfunction further propagates features of AD, such as lipid droplet accumulation, increased amyloid burden, and neuroinflammation, to name a few (67). Notably, while we consider ApoE4 and HSV-1 to be independent risk factors, since both impact the function of microglia, there is likely a synergistic impact on microglial dysfunction that preceeds AD pathology. Indeed, epidemiological studies report a synergistic risk between ApoE4, HSV-1 infection, and AD risk (68). Further studies investigating the interaction between ApoE4, HSV-1 infection, and AD risk are warranted.

Understanding regional consequences of HSV-1 reactivation in the specific brain region could point to an array of different neurological complications (69). The brainstem has many autonomic and sensory processes, including many reticular nuclei. Many basic motor functions include coordination and reflexes (70). For example, the medullary reticular nucleus is well known for modulation of nociception to acute inflammatory pain; loss of inhibition to the reticular nuclei can lead to increased response to pain (71). Indeed, herpesvirus infection is associated with many chronic pain disorders, including post-herpetic neuralgia and trigeminal neuralgia (72, 73) (74).

Neuropsychiatric disorders have also been associated with herpes virus infection, including anxiety (75, 76). Multiple regions were identified in this study to be infected with HSV-1 that are involved in anxiety and sleep/wake cycles, including the noradrenergic center (LC) and hypothalamus (77, 78). It is important to note that the LC and hypothalamus are known areas to be dysregulated in AD and are associated with non-cognitive deficits (46, 79–84). Our results point to vulnerable points of entry to the CNS that may accelerate non-cognitive deficits seen in neurodegenerative disorders (85).

Lastly, we did not find HSV-1 in the hippocampus or the cortex. Yet, those regions are involved in memory and higher-order executive functions, which are altered in patients with herpes simplex encephalitis (69). While these regions may become infected following multiple reactivation events, as has been shown by other groups, we emphasize that this mouse model is suitable for investigating HSV-1 brain entry and spread apart from mimicking herpes simplex encephalitis. Together, these findings expand our understanding of the nuance pattern of HSV-1 infection within the CNS, offering insights into different neurological complications and pathology associated with these distinct brain regions that become infected with HSV-1.

## Acknowledgments

This research was supported by grants from the National Institute of Aging R01AG079193 (D.R.; M.A.N.), the Department of Otolaryngology at the University of Colorado Anschutz Medical Campus through the National Institute of Deafness and Other Communication Disorders 2T32DC012280-06A1 (C.S.N.), and NIH/NCATS Colorado CTSA K12 TR004412 (C.S.N.).

