## Supplementary figures and images for "Olfactory and Trigeminal Routes of HSV-1 CNS Infection with Regional Microglial Heterogeneity"

### Supplemental Figure 1

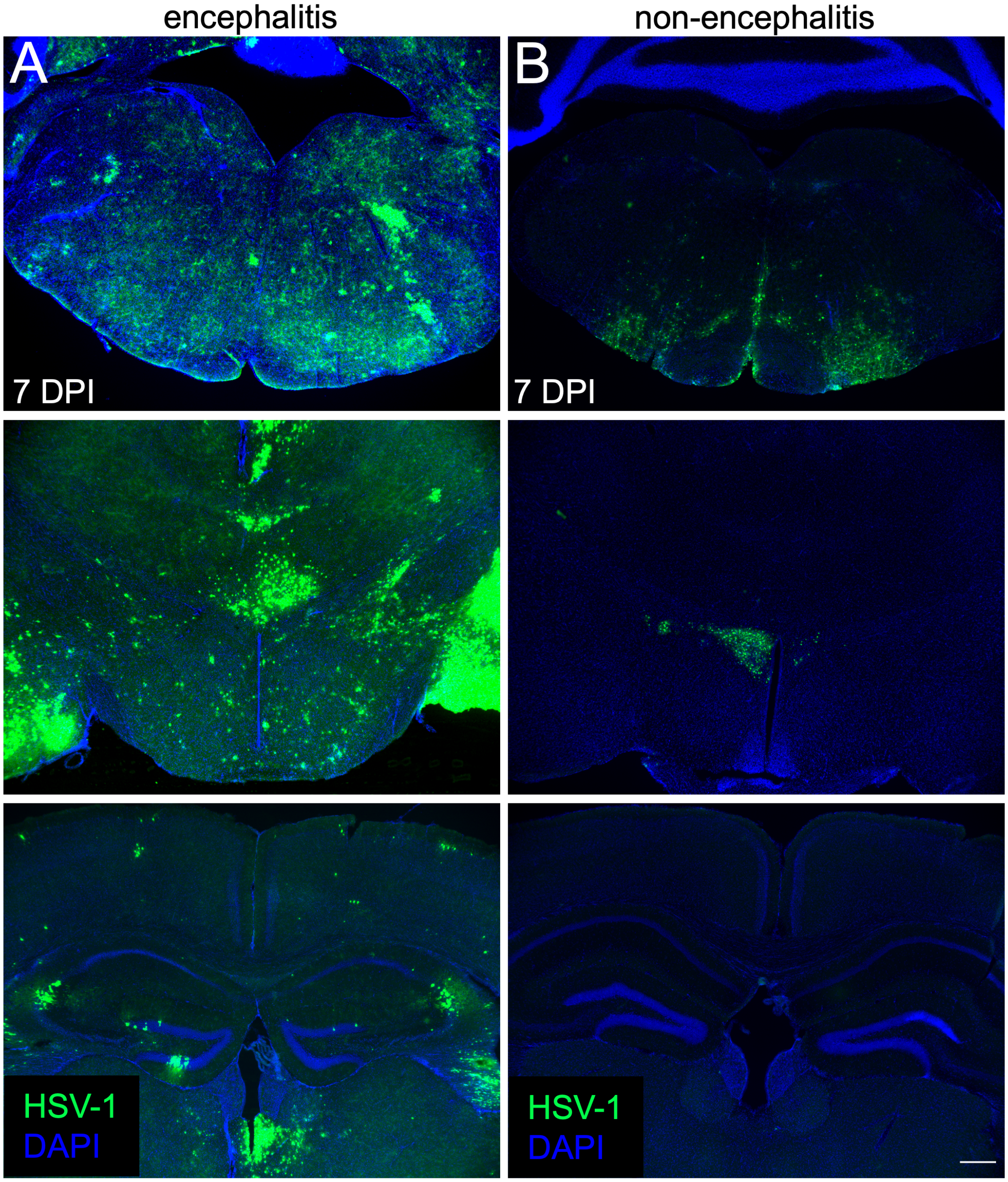

### Supplemental Figure 3

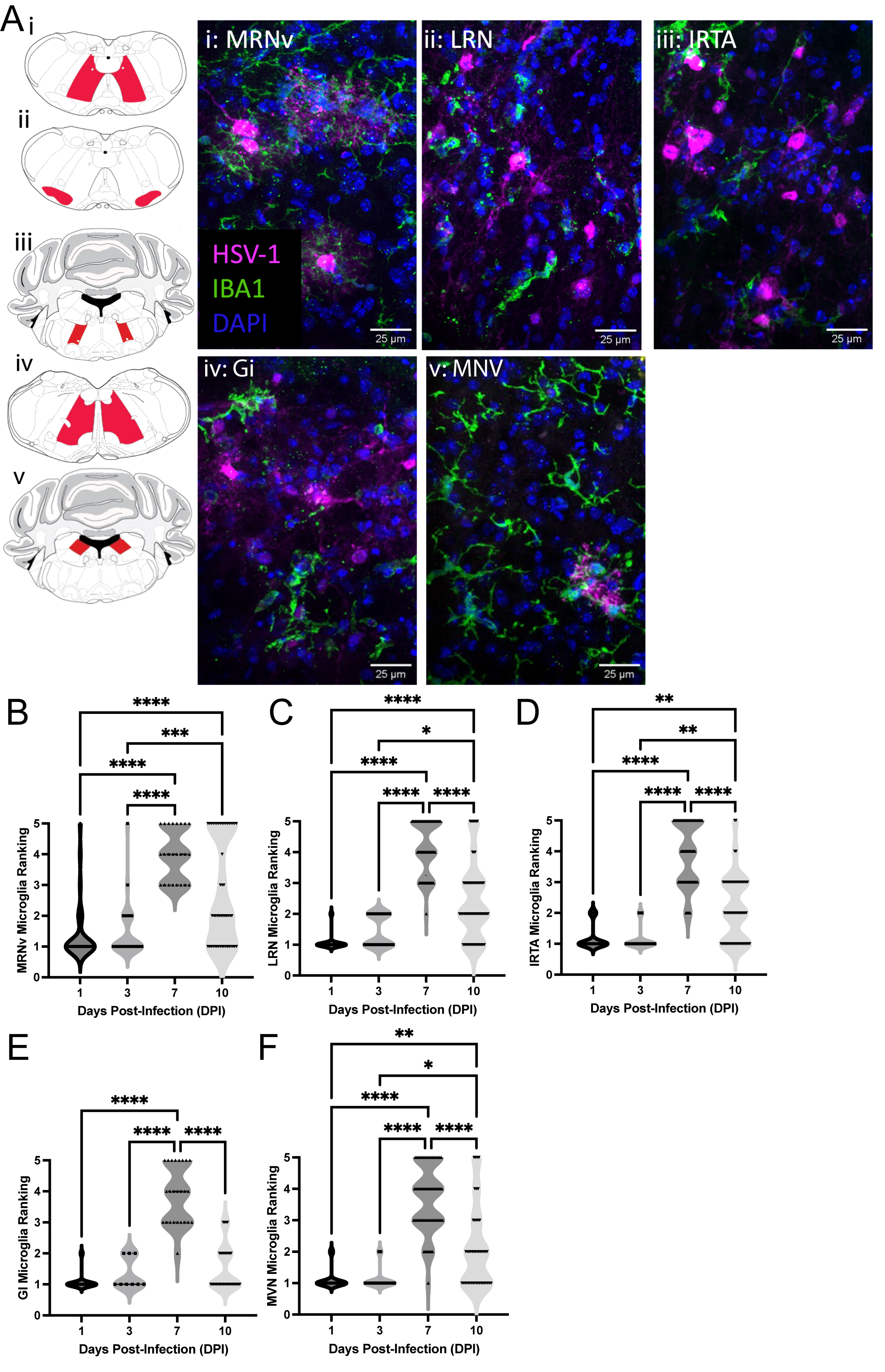
