## Supplemental Figure 2 for "Olfactory and Trigeminal Routes of HSV-1 CNS Infection with Regional Microglial Heterogeneity"

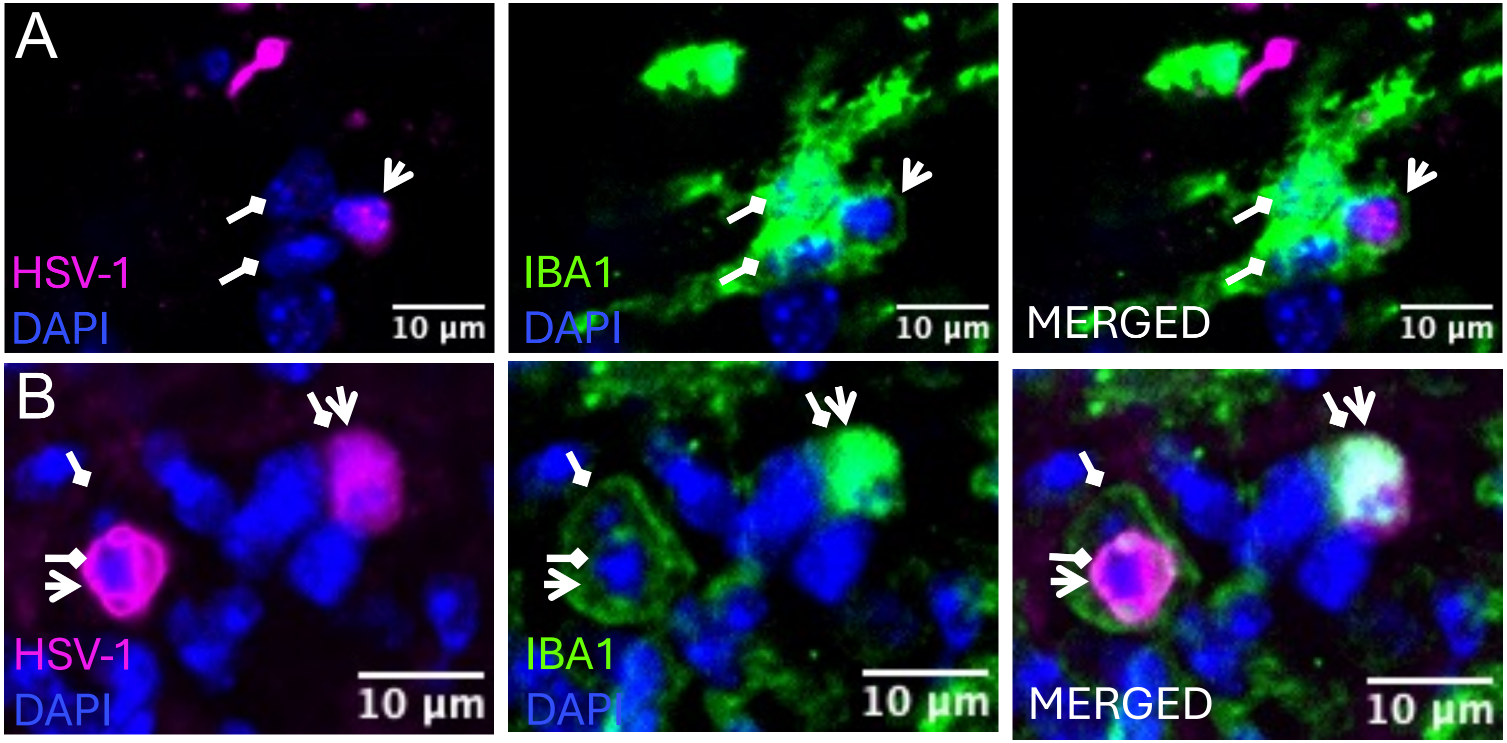


Supplemental Fig. 2. Colocalization of IBA1 and HSV-1. Single plane images ( $z = 0.34\mu$ m) were taken in the brainstem of HSV-1 infected C57Bl/6 mice at 7 days post-infection. (A) two IBA-1 positive cells engulfing an HSV-1 positive IBA1-negative cells. (B) Two IBA1-positive cells colocalizing with HSV-1 antigen (right). Diamond arrow: IBA1+ cells, Arrow: HSV-1 infected cell.
